# Potent, specific MEPicides for treatment of zoonotic staphylococci

**DOI:** 10.1101/626325

**Authors:** Rachel L. Edwards, Isabel Heueck, Soon Goo Lee, Ishaan T. Shah, Andrew J. Jezewski, Justin J. Miller, Marwa O. Mikati, Xu Wang, Robert C. Brothers, Kenneth M. Heidel, Carey-Ann D. Burnham, Sophie Alvarez, Stephanie A. Fritz, Cynthia S. Dowd, Joseph M. Jez, Audrey R. Odom John

**Author notes:** Correspondence and requests for materials should be addressed to R.L.E. or A.R.O.J. Isabel Heueck, Sandoz Deutschland/Hexal AG, Oberhaching, Germany; Xu Wang, Brown University Providence, RI, USA; Soon Goo Lee, University of North Carolina-Wilmington, Wilmington, NC, USA; Robert C. Brothers, Naval Surface Warfare Center, Indian Head, MD, USA.

## Abstract

Coagulase-positive staphylococci, which frequently colonize the mucosal surfaces of animals, also cause a spectrum of opportunistic infections including skin and soft tissue infections, urinary tract infections, pneumonia, and bacteremia. However, recent advances in bacterial identification have revealed that these common veterinary pathogens are in fact, zoonoses that cause serious infections in human patients. The global spread of multidrug-resistant zoonotic staphylococci, in particular the emergence of methicillin-resistant organisms, is now a serious threat to both animal and human welfare. Accordingly, new therapeutic targets that can be exploited to combat staphylococcal infections are urgently needed. Enzymes of the methylerythritol phosphate pathway (MEP) of isoprenoid biosynthesis represent potential targets for treating zoonotic staphylococci. Here we demonstrate that fosmidomycin (FSM) inhibits the first step of the isoprenoid biosynthetic pathway catalyzed by deoxyxylulose phosphate reductoisomerase (DXR) in staphylococci. In addition, we have both enzymatically and structurally determined the mechanism by which FSM elicits its effect. Using a forward genetic screen, the glycerol-3-phosphate transporter GlpT that facilitates FSM uptake was identified in two zoonotic staphylococci, *Staphylococcus schleiferi* and *Staphylococcus pseudintermedius*. A series of lipophilic ester prodrugs (termed MEPicides) structurally related to FSM were synthesized, and data indicate that the presence of the prodrug moiety not only substantially increased potency of the inhibitors against staphylococci, but also bypassed the need for GlpT-mediated cellular transport. Collectively, our data indicate that the prodrug MEPicides selectively and robustly inhibit DXR in zoonotic staphylococci, and further, DXR represents a promising, druggable target for future development.

**Author Summary:** The proliferation of microbial pathogens resistant to the current pool of antibiotics is a major threat to public health. Drug resistance is pervasive in staphylococci, including several species that can cause serious zoonotic infections in humans. Thus, new antimicrobial agents are urgently need to combat these life-threatening, resistant infections. Here we establish the MEP pathway as a promising new target against zoonotic staphylococci. We determine that fosmidomycin (FSM) selectively targets the isoprenoid biosynthesis pathway in zoonotic staphylococci, and use forward genetics to identify the transporter that facilitates phosphonate antibiotic uptake. Employing this knowledge, we synthesized a series of potent antibacterial prodrugs that circumvent the transporter. Together, these novel prodrug inhibitors represent promising leads for further drug development against zoonotic staphylococci.

## Introduction

Coagulase-positive staphylococci, such as *S. pseudintermedius* and *S. schleiferi* subsp. *coagulans*, are leading causes of skin, soft tissue, and invasive infections in companion animals such as dogs and cats. In addition, these organisms cause zoonotic infections in humans that are clinically indistinguishable from infections with *S. aureus* including pneumonia, skin and soft tissue infections, hardware infections, and bacteremia(1–5). Newer clinical microbiological techniques, such as mass spectrometry, now readily distinguish *S. aureus* from zoonotic coagulase-positive staphylococci, which were previously often misidentified(3,6,7). Thus, there is a growing recognition of the importance of zoonotic staphylococci in human disease. Because *mecA*-mediated methicillin resistance is on the rise in both veterinary and human clinical isolates, new antibacterial strategies to specifically target zoonotic staphylococci are highly desirable(8–10).

Two distinct and independent pathways for isoprenoid biosynthesis have evolved, the mevalonate pathway and a mevalonate-independent route that proceeds through methylerythritol phosphate, called the MEP pathway(11). Unusual among bacteria, the least common ancestor of all *Staphylococcus* spp. appears to have possessed both pathways. Primate-associated staphylococcal lineages, including *S. aureus*, possess the mevalonate pathway, and evidence suggests that mevalonate pathway activity is required for peptidoglycan synthesis, growth, and virulence(12–14). In contrast, nonprimate-associated staphylococcal species, including *S. pseudintermedius* and *S. schleiferi*, utilize the MEP pathway for isoprenoid biosynthesis(15). Importantly, humans and other mammals lack homologs of MEP pathway enzymes, and MEP pathway activity is required for cellular growth in all organisms in which it has been experimentally determined(16–21). Thus, new chemical inhibitors of MEP pathway enzymes hold promise as effective antimicrobials that may provide a high margin of safety.

The first dedicated enzyme of the MEP pathway, deoxyxylulose phosphate reductoisomerase (DXR; E.C. 1.1.1.267), is rate-limiting for MEP pathway activity. DXR is known to be susceptible to small molecule inhibition. For example, the phosphonic acid antibiotic fosmidomycin (FSM) is a slow, tight-binding, competitive inhibitor of DXR(22). FSM is safe and well-tolerated in humans and animals(23–25). Unfortunately, FSM has poor oral bioavailability and a short serum half-life, which has hampered clinical efficacy. Moreover, the charged nature of FSM and its phosphonate analogs has challenged their clinical development as the compounds are excluded from cells unless actively transported. As a result, many microorganisms, such as *Mycobacterium tuberculosis* and *Toxoplasma gondii*, are inherently resistant to FSM (due to poor cellular uptake) even though FSM potently inhibits their DXR orthologs *in vitro*(16,18,26). In Gram-negative organisms, FSM import is dependent on a glycerol-3-phosphate/Pi antiporter (GlpT), and FSM resistance can be achieved by reduced expression or activity of GlpT(27,28).

In this work, we use the highly specific inhibitor FSM to chemically validate the MEP pathway enzyme DXR as an essential, druggable antibacterial target for zoonotic staphylococcal infections. Furthermore, we establish the structural and enzymatic mechanism of staphylococcal DXR inhibition by FSM. Using a chemical genomics approach, we define the genetic basis of FSM resistance in zoonotic staphylococci and define the FSM transporter GlpT in these strains. Finally, we reveal that structurally related lipophilic ester prodrugs (called MEPicides) yield substantially increased potency and circumvent the need for GlpT-dependent import. Thus, lipophilic prodrugs provide a promising new approach to combat zoonotic staphylococcal infections.

## Results

### Anti-staphylococcal activity of canonical MEP pathway inhibitors

Because previous evidence had suggested that zoonotic staphylococci might be sensitive to MEP pathway inhibition, we quantified the dose-dependent antibacterial effects of FSM and FR-900098, a structurally similar DXR inhibitor (Table 1)(15). FSM was 5-10-fold more potent against both *S. schleiferi* (IC_50_ = 0.78 ± 0.13 μM) and *S. pseudintermedius* (IC_50_ = 0.31 ± 0.04 μM), respectively (Table 1), despite modest chemical differences between the two inhibitors. Data indicate that both compounds elicit their effect via a bacteriostatic mechanism-of-action, as neither caused a substantial drop in viable cells (Fig S1). Because *S. aureus* does not utilize the MEP pathway for isoprenoid biosynthesis, neither FSM nor FR-900098 inhibit *S. aureus* growth (Table 1). Together, these data indicate that both *S. schleiferi* and *S. pseudintermedius* have a functional MEP pathway that is required for bacterial growth.

**Table 1.**
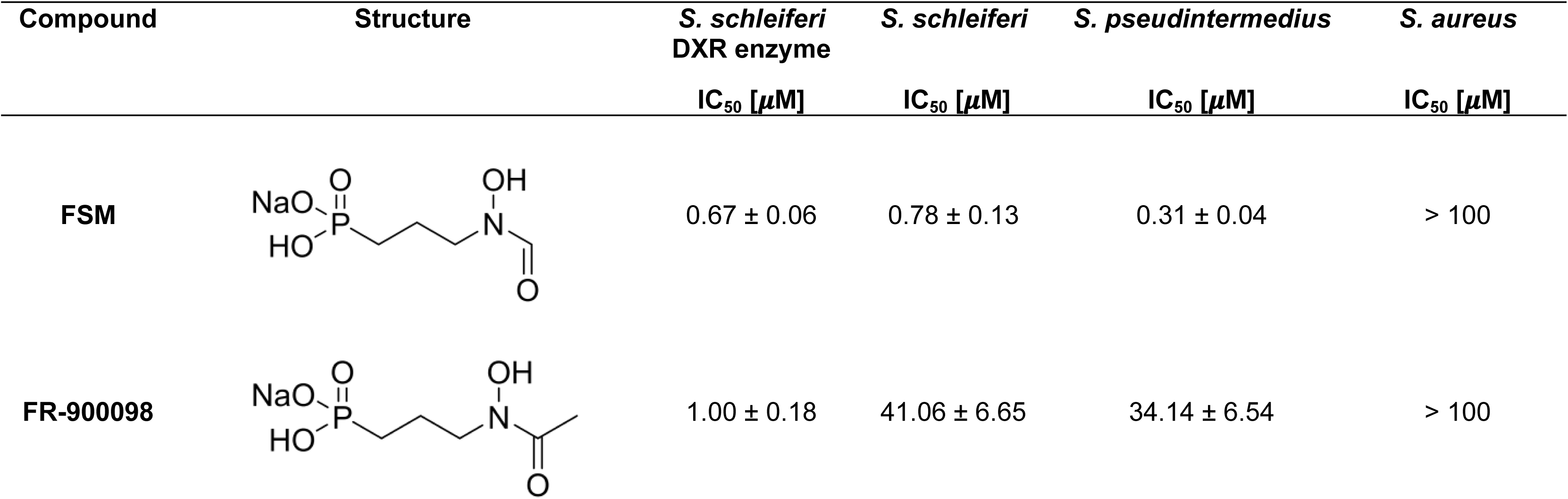
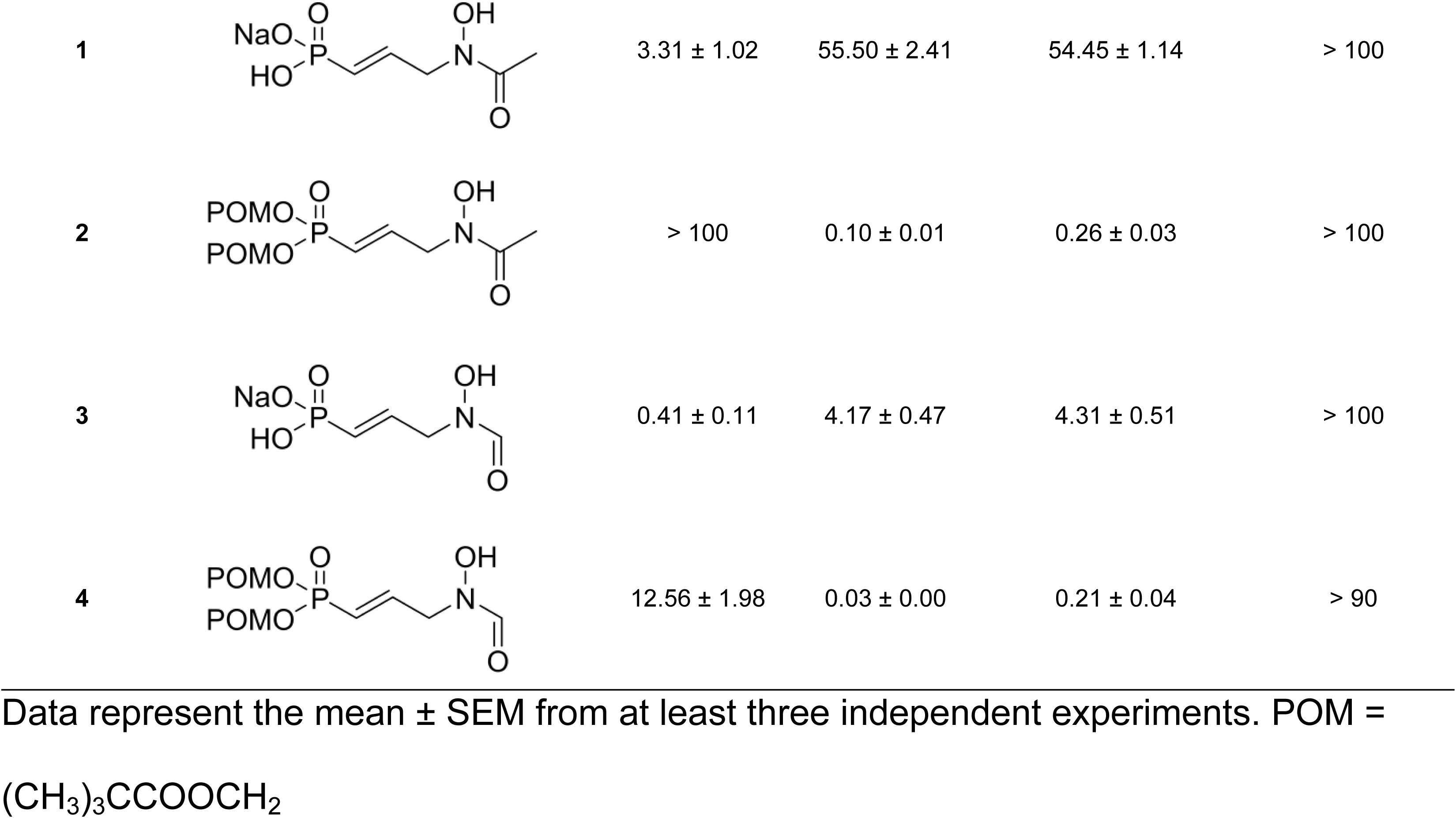
Inhibitory effect of MEPicides against the *S. schleiferi* DXR enzyme and *in vitro* activity against Staphylococcus spp.

| Compound | Structure | *S. schleiferi*<br>DXR enzyme<br>$IC_{50} [\mu M]$ | *S. schleiferi*<br>$IC_{50} [\mu M]$ | *S. pseudintermedius*<br>$IC_{50} [\mu M]$ | *S. aureus*<br>$IC_{50} [\mu M]$ |
| --- | --- | --- | --- | --- | --- |
| FSM | | $0.67 \pm 0.06$ | $0.78 \pm 0.13$ | $0.31 \pm 0.04$ | > 100 |
| FR-900098 | | $1.00 \pm 0.18$ | $41.06 \pm 6.65$ | $34.14 \pm 6.54$ | > 100 |
| 1 | | $3.31 \pm 1.02$ | $55.50 \pm 2.41$ | $54.45 \pm 1.14$ | > 100 |
| 2 | | > 100 | $0.10 \pm 0.01$ | $0.26 \pm 0.03$ | > 100 |
| 3 | | $0.41 \pm 0.11$ | $4.17 \pm 0.47$ | $4.31 \pm 0.51$ | > 100 |
| 4 | | $12.56 \pm 1.98$ | $0.03 \pm 0.00$ | $0.21 \pm 0.04$ | > 90 |

Data represent the mean ± SEM from at least three independent experiments. POM = $(\text{CH}_3)_3\text{CCOOCH}_2$

### Fosmidomycin inhibits isoprenoid metabolism in zoonotic staphylococci

To establish the presence of MEP pathway intermediates and to determine the cellular mechanism-of-action of FSM, we performed targeted metabolic profiling of MEP pathway intermediates in *S. schleiferi* and *S. pseudintermedius*, with and without drug treatment. We confirmed that both species contain MEP pathway intermediates, including the DXR substrate, deoxyxylulose 5-phosphate (DOXP), and the downstream metabolite, methylerythritol cyclodiphosphate (MEcPP) (Fig 1). Upon FSM treatment, intracellular levels of DOXP increase dramatically (23.8-fold; p < 0.05 and 34.8-fold; p < 0.05 for *S. schleiferi* and *S. pseudintermedius*, respectively), consistent with DXR inhibition. Similarly, intracellular levels of MEcPP are substantially reduced following FSM treatment (4.5-fold; p < 0.01 and 2.4-fold; p < 0.05 for *S. schleiferi* and *S. pseudintermedius*, respectively), consistent with FSM-mediated reduction in MEP pathway metabolism. Together, these data confirm the presence of active MEP pathway metabolism in zoonotic staphylococci and establish that FSM inhibits growth through MEP pathway inhibition.

**Fig 1.**
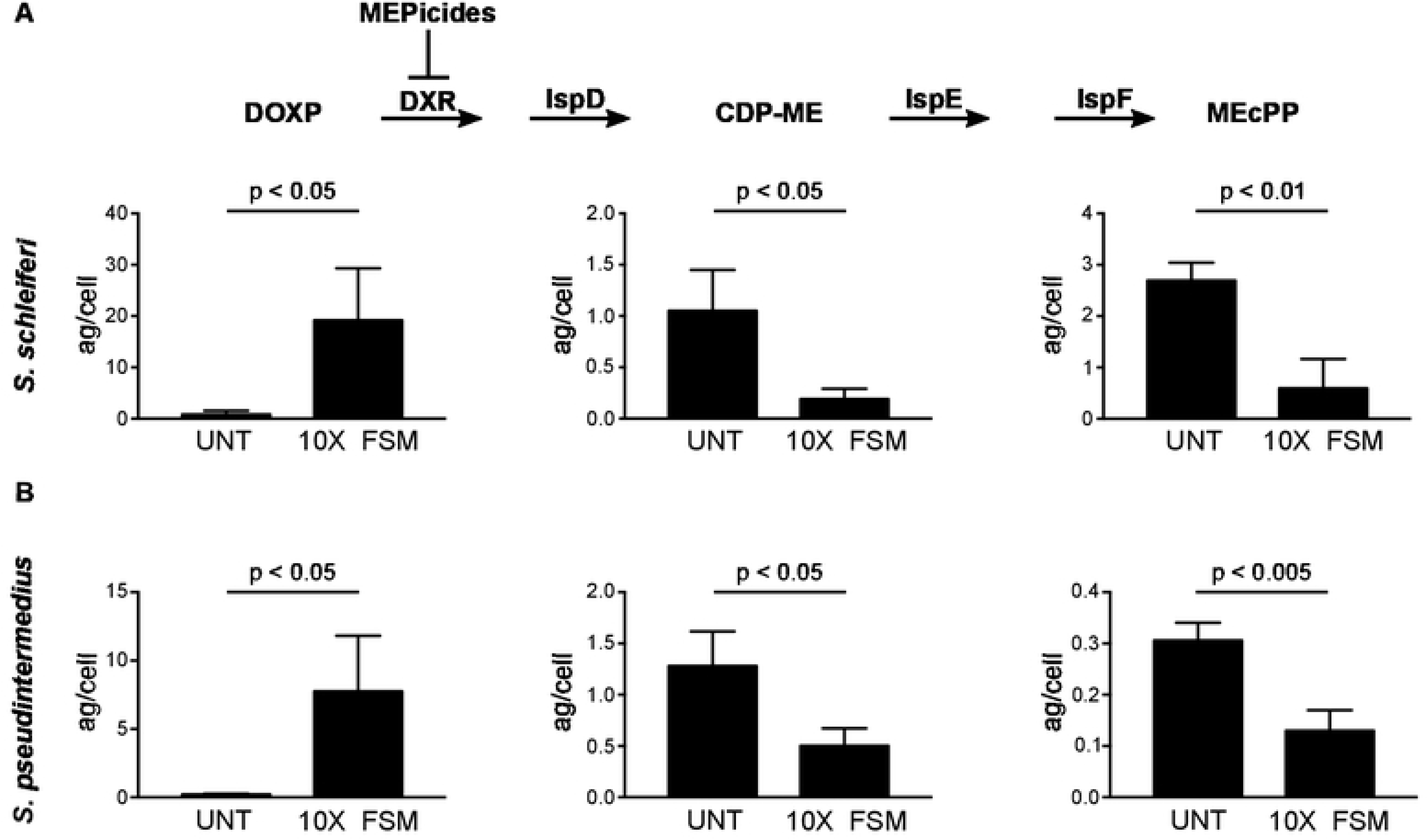
FSM inhibits the MEP pathway in *Staphylococcus* spp. MEP pathway metabolites were compared between untreated (UNT) *S. schleiferi* (A) and *S. pseudintermedius* (B) and bacteria treated with FSM at 10x the respective IC_50_ values. After 2 h treatment, bacterial cells were harvested and the cell pellets analyzed by LC-MS/MS. Displayed are the means ± SD of the metabolite levels from three independent experiments. P-values were determined using a Student’s *t*-test.

### Fosmidomycin is a competitive inhibitor of *S. schleiferi* DXR

To establish the enzymatic mechanism-of-action of DXR inhibitors against staphylococci, we cloned and purified *S. schleiferi* DXR (Fig S2; Table S1). Enzymatic characterization of DXR confirmed a Michaelis constant (K_m_) [DOXP] (0.52 ± 0.08 mM), similar to that of other DXR orthologs (Fig 2A)(29,30). Both FSM and FR-900098 inhibit *S. schleiferi* DXR in a dose-dependent manner (Table 1). Further, we confirm that DXR inhibition by FSM is competitive with respect to the DOXP substrate, with a K_i_ [DOXP] of 0.29 ± 0.022 μM (Fig 2B).

**Fig 2.**
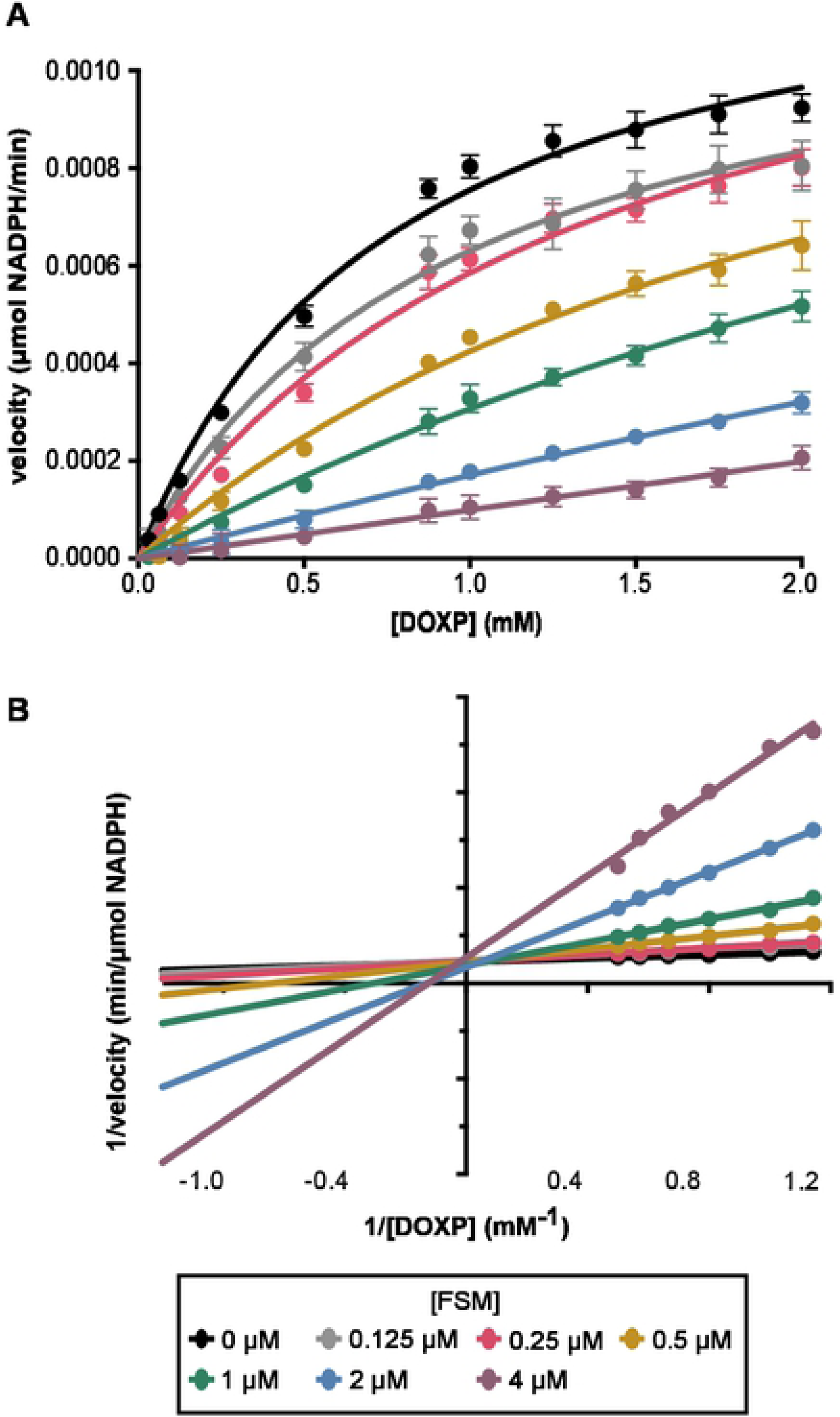
Inhibition of staphylococcal DXR by FSM is competitive with DOXP. (A) *S. schleiferi* DXR velocity in μmol NADPH/min with respect to the DOXP concentration in mM. Displayed are the means ± SD from three independent experiments. (B) Lineweaver-Burk double reciprocal plots of *S. schleiferi* DXR activity over a range of DOXP substrate concentrations, for illustrative purposes only.

### Structural basis of fosmidomycin inhibition

To establish the structural basis of FSM action, we solved the three-dimensional structures of *S. schleiferi* DXR as an apoenzyme and a FSM complex to 2.15 Å and 2.89 Å resolution, respectively (Table S2; Fig 3). *S. schleiferi* DXR is a physiologic dimer with each monomer related by crystallographic symmetry (Fig 3A). A DALI search identified multiple DXR from *Escherichia coli, Plasmodium falciparum, M. tuberculosis*, and other microbes (Z-scores: 49-51; r.m.s.d. ∼1.6 Å^2^ for 370-400 C_α_-atoms; 39-40% amino acid sequence identity)(31–36). The monomer consists of three regions (Fig 3A): an N-terminal α/β-domain with a central 7-stranded β-sheet (β1-β7) and 7 α-helices that serves as the nucleotide binding site; a middle region of the protein that includes a second β-sheet (β8-β11) and 4 α-helices (α8 and α12-α14); and a C-terminal α-helical domain (α9-α11 and α15-α18) that locks FSM into the active site(37).

**Fig 3.**
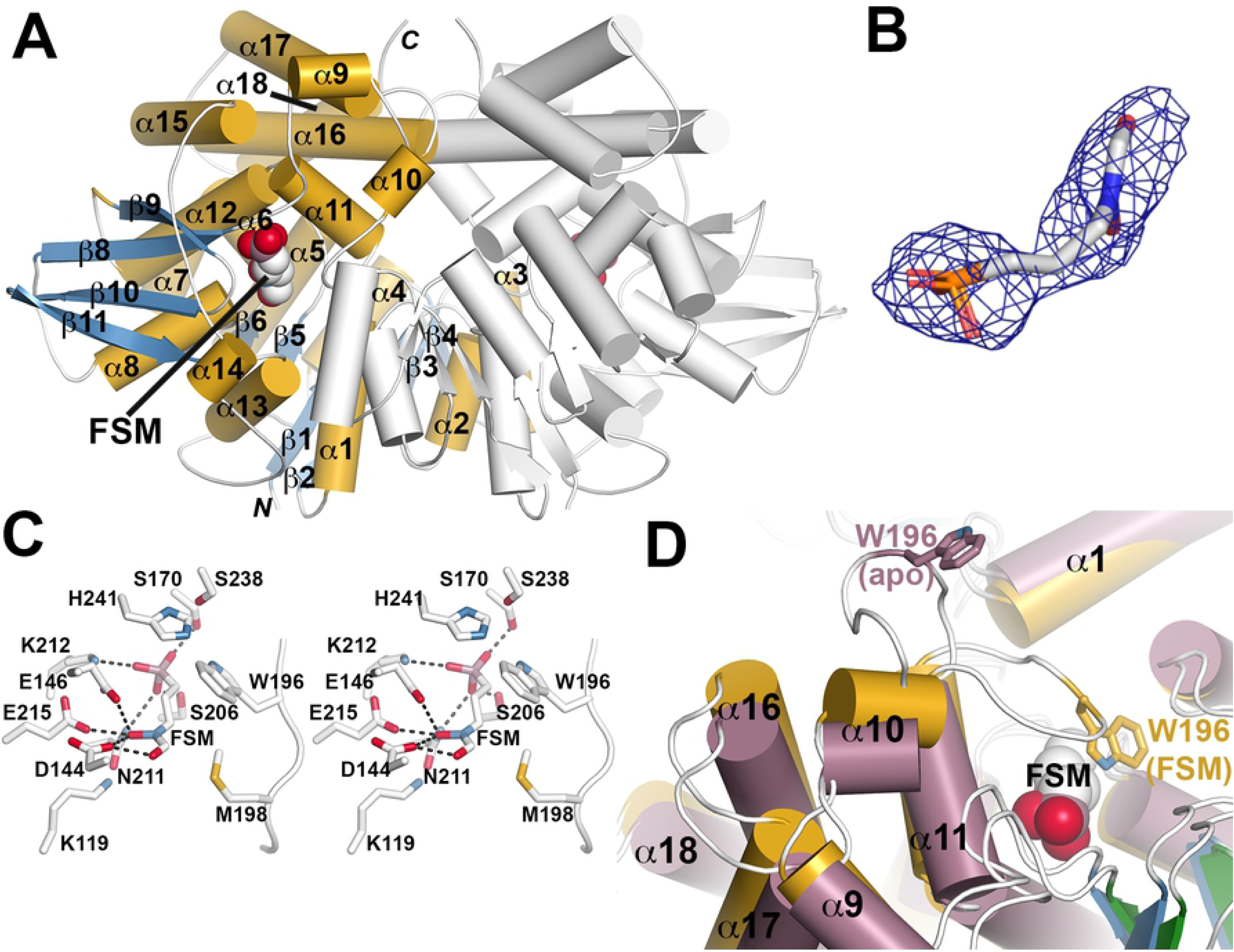
Crystal structure of *S. schleiferi* DXR. (A) Overall structure of the *S. schleiferi* DXR•FSM complex. The dimer is shown as a ribbon diagram with α-helices and β-strands of one monomer colored gold and blue, respectively. The position of FSM (space-filling model) in one monomer is indicated. (B) Electron density for FSM is shown as a 2F_o_-F_c_ omit map (1 σ). (C) Stereoview of FSM binding in the active site. Dotted lines indicate protein-ligand interactions. (D) Comparison of *S. schleiferi* DXR apoenzyme and FSM complex structures. Structural changes in the active site region between the apoenzyme (rose) and FSM complex (gold) are shown. The major change in the position of the α10-α11 loop is emphasized by the position of Trp196 in each structure.

Clear electron density for FSM was observed in the active site (Fig 3B) and revealed multiple protein-ligand interactions (Fig 3C). Interactions with Ser170, Ser206, Asn211, and Lys212 positions the FSM phosphonate toward the catalytic histidine (His241) and the NADP(H) binding site. The hydroxamic acid of the ligand contacts Asp144, Glu146, and Glu215. Additional van der Waals contacts are provided by Trp196, which resides in the α10-α11 loop. Comparison of the *S. schleiferi* DXR apoenzyme and FSM complex structures reveals how the C-terminal capping region (α9-α11 and α16-18) shift position to allow for the α10-α11 loop to position Trp196 adjacent to the inhibitor (Fig 3D). Movement of this flexible loop is a key feature for FSM inhibition of DXR from a variety of microorganisms(38). The residues that interact with FSM in the *S. schleiferi* DXR are conserved in the crystal structures of DXR from *E. coli, P. falciparum*, and *M. tuberculosis* with some variation in the sequence of the α10-α11 loop, although the tryptophan that contacts FSM is conserved in all these enzymes(34,36,37).

### Resistance selection reveals a candidate FSM transporter, GlpT

To establish the molecular basis of compound uptake, we performed independent, parallel, forward genetic screens for FSM resistance in both *S. schleiferi* and *S. pseudintermedius* (Fig 4A). Candidate FSM resistant (FSM^R^) strains were colony purified and resistance was quantified by MIC determination (Fig 4B and Table S3). For both *S. schleiferi* and *S. pseudintermedius*, FSM^R^ strains possessed FSM MICs >100-fold higher than the wild-type parental lines. We employed whole genome sequencing to characterize the single-nucleotide polymorphisms (SNPs) that were present in the resistant strains (Table S4). In both species, FSM selective pressure enriched for new nonsynonymous changes in a single homologous locus, RN70_03745 (10/11 *S. schleiferi* strains) and SPSE_0697 (10/12 *S. pseudintermedius* strains) (Figs. S3A and S3B). These loci are close homologs (>90% sequence identity and 95.4% sequence similarity), which belong to the glycerol-3-phosphate transporter (GlpT) subfamily (Interpro: IPR005267) of the major facilitator superfamily (MFS) family of proteins (Interpro: IPR011701). These data suggest a model in which GlpT mediates FSM import, such that loss of GlpT function confers FSM resistance.

**Fig 4.**
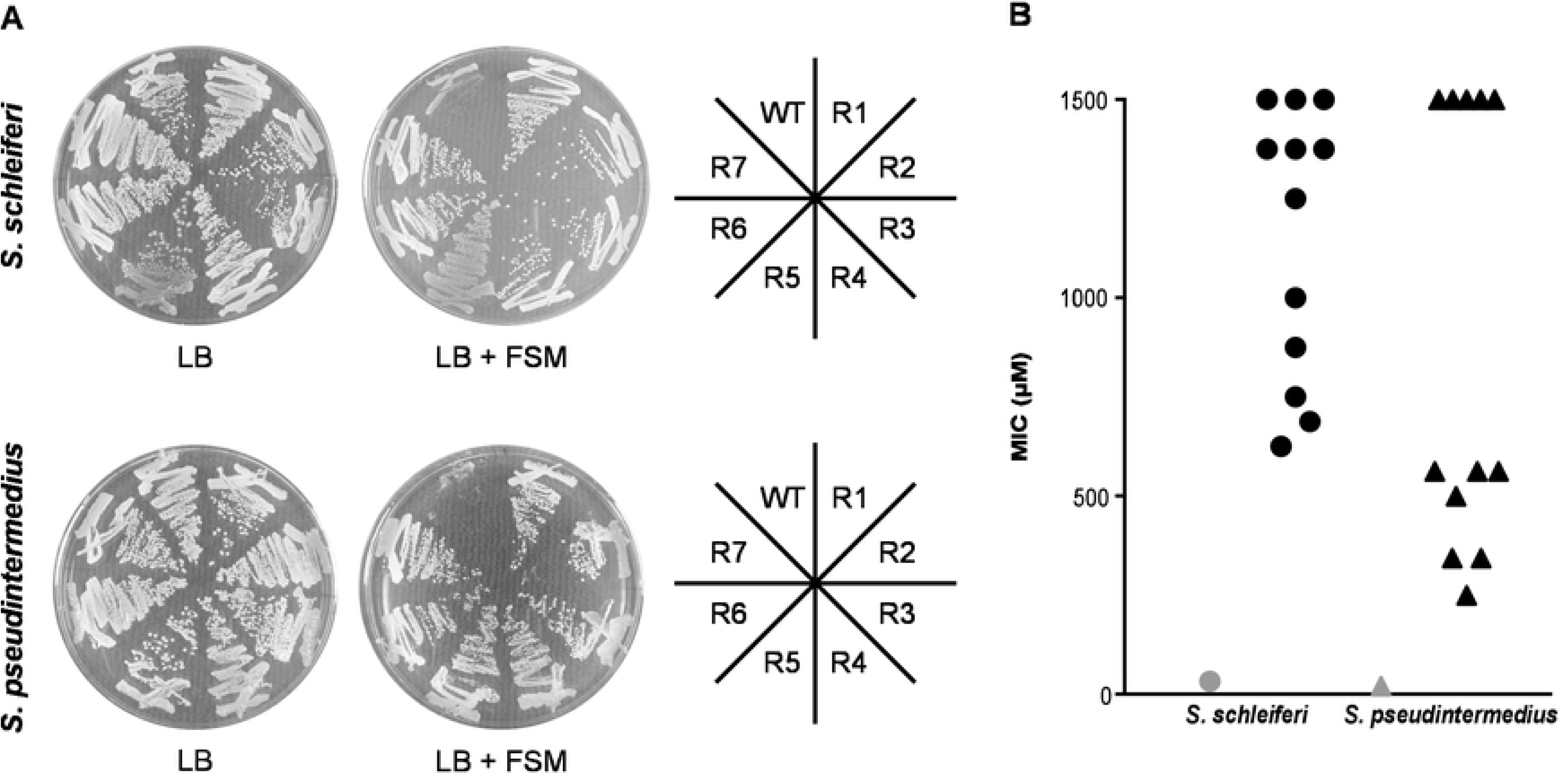
Successful evolution of FSM resistance. (A) Wild-type and FSM-resistant isolates from *S. schleiferi* (top) or *S. pseudintermedius* (bottom) were struck on LB agar plates with (right) and without (left) 32 μM FSM. (B) Distribution of the MIC values for WT (gray) and FSM-resistant mutants (black) from *S. schleiferi* (circles) and *S. pseudintermedius* (triangles). Displayed are the mean values for each strain from three independent experiments.

### Fosmidomycin-resistance alleles of the candidate transporter, GlpT

We predicted that the FSM-resistance alleles likely reduce GlpT function. In *S. schleiferi*, nine distinct alleles were found with GlpT changes: two with nonsense mutations and seven others with amino acid variants that are predicted to be highly deleterious (Polyphen-2 score >0.9; Table S3)(39). Similarly, in *S. pseudintermedius*, a total of seven distinct alleles were identified with GlpT sequence changes. Of these, one contained a nonsense mutation and six other GlpT variants contained amino acid substitutions that are strongly predicted to reduce function (Polyphen-2 score >0.9; Table S3). FSM-resistant variants map along the length of the nearly 50 Kd GlpT transporter, in both *S. schleiferi* and *S. pseudintermedius* (Figs S3A and S3B). Altogether, the finding of multiple independent loss-of-function alleles, including nonsense mutations, in two different selections in distinct organisms, strongly suggests that reduced GlpT function is responsible for FSM resistance in these strains.

### Lipophilic ester prodrugs with improved anti-staphylococcal potency

Due to their charged nature, phosphonic acid antibiotics have poor cellular penetration and bioavailability, and serum half-lives are relatively brief(23,25,40). In the ongoing effort to develop new treatments for malaria and tuberculosis by improving upon the drug-like properties of phosphonates, numerous lipophilic ester prodrugs that target DXR have been generated(41–53) Our phosphonate parent compounds (1 and 3) are similar in anti-staphylococcal potency to FSM and FR-900098 (Table 1); however, lipophilic modification of either compound dramatically improves potency (in most cases by 100-fold) against both *S. schleiferi* and *S. pseudintermedius* (compare compound 1 to its prodrug, compound 2, and compound 3 to its prodrug, compound 4) (Table 1). As expected, prodrugs 2 and 4 poorly inhibit purified recombinant *S. schleiferi* DXR *in vitro*, since cleavage of the prodrug moiety is required for activity (Table 1). Our data suggest that lipophilic ester modifications improves uptake of the DXR inhibitors, and that active phosphonates are released intracellularly for target inhibition (model, Fig 6).

### Lipophilic prodrugs bypass need for GlpT-mediated transport

We anticipated that our lipophilic ester prodrugs do not require active cellular transport. To evaluate whether GlpT is required for prodrug uptake, we characterized the MEPicide sensitivity of four different FSM^R^ *glpT* mutant *S. schleiferi* strains. As expected, we find that FSM^R^ *glpT* strains are cross-resistant to the phosphonate parent drug (compound 3), suggesting a common mechanism of transport (Fig 5). In contrast, FSM^R^ *glpT* strains remain sensitive to the MEPicide prodrugs compounds 2 and 4, supporting a model in which GlpT mediates phosphonate transport, with the ester modifications substantially improving cellular uptake (Fig 6)(21).

**Fig 5.**
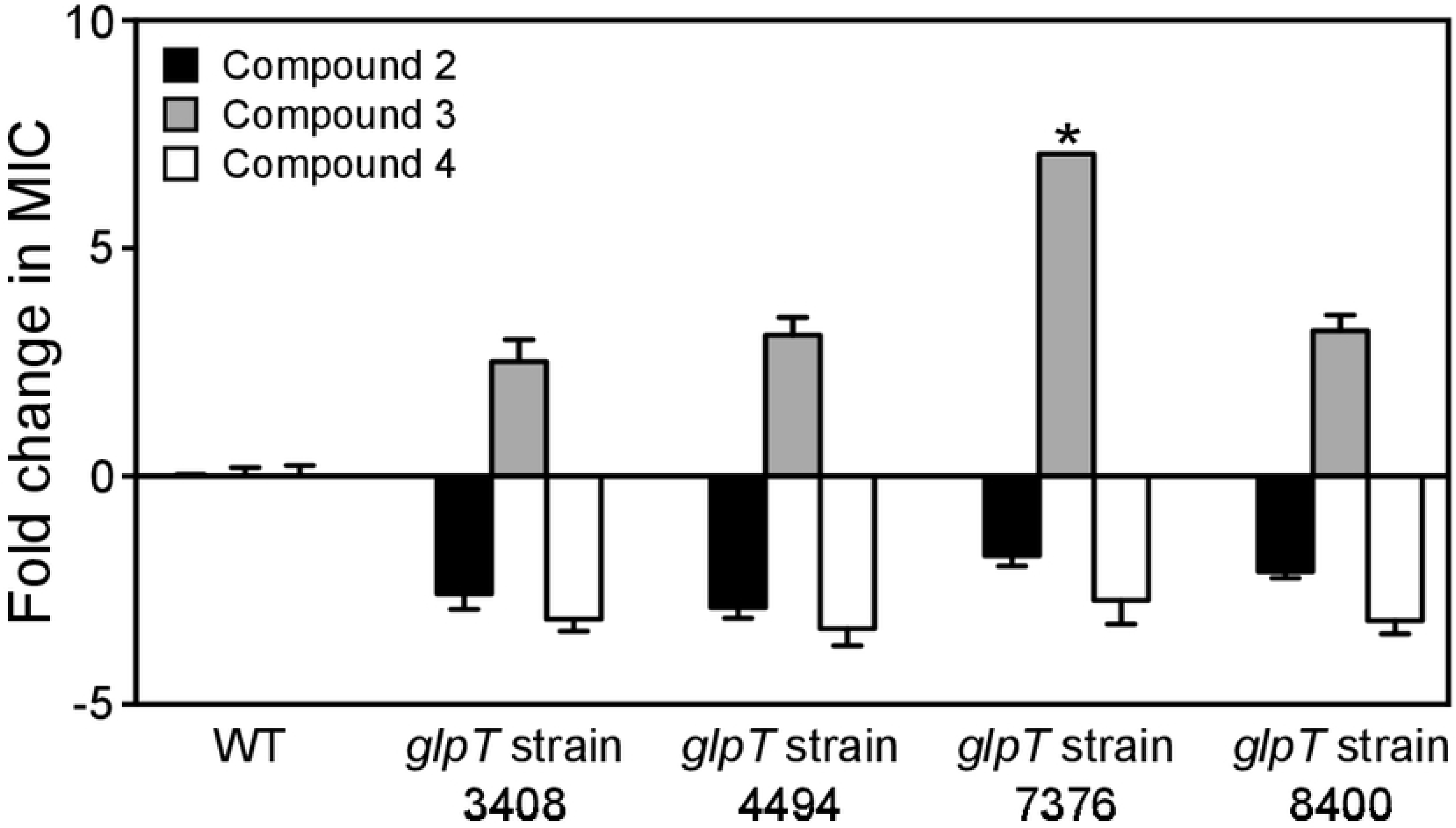
*glpT* mutant staphylococci are sensitive to MEPicide prodrugs. Wild-type (WT) and FSM-resistant, *glpT* mutant *S. schleiferi* isolates (strains 3408, 4494, 7376, and 8400) were treated with MEPicides and the MIC values determined during overnight growth. Displayed are the mean values of the fold change (resistant isolate/WT) ± SEM from at least three independent experiments. *MIC values observed for *glpT* strain 7376 were identical in three independent experiments performed in technical duplicate.

**Fig 6.**
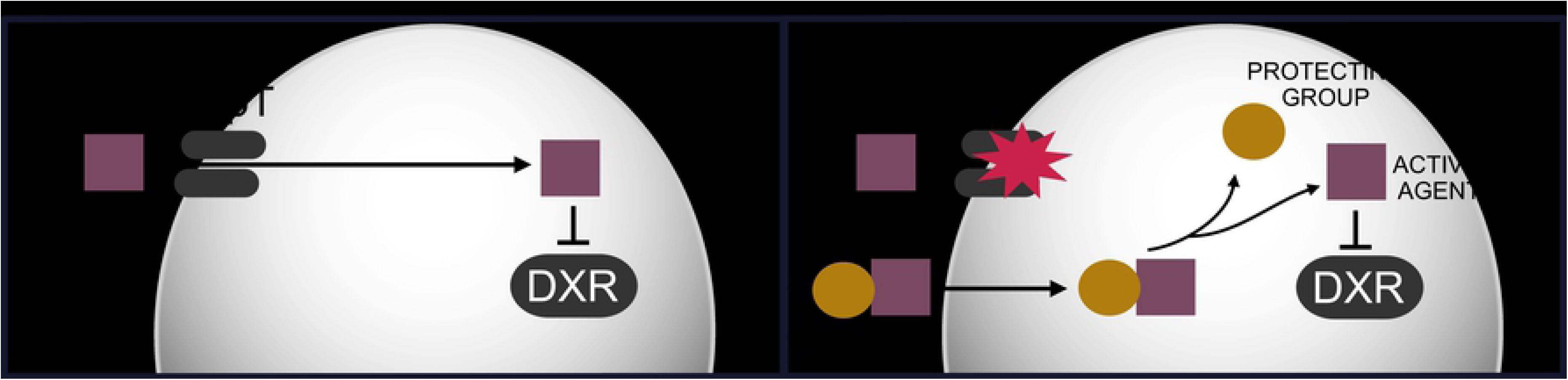
Model. In wild-type zoonotic staphylococci, GlpT transports the MEP pathway inhibitor FSM intracellularly where it inhibits its target, DXR. In staphylococci with *glpT* mutations, FSM is excluded from cells, resulting in FSM resistance. In contrast, lipophilic prodrug MEPicides do not require active transport and remain effective.

## Discussion

*S. schleiferi* and the *Staphylococcus intermedius* group (SIG) (including *S. pseudintermedius, S. intermedius*, and *S. delphini)* cause pyodermic infections in companion animals, such as dogs and cats(8). Treatment of these infections is complicated by rising rates of antimicrobial resistance, particularly methicillin-resistance(54). A growing recognition that SIG species also cause zoonotic human infections, indistinguishable from infections with *S. aureus*, has led to new urgency in the search for additional therapeutics against these organisms. The non-mevalonate pathway of isoprenoid biosynthesis through MEP has been previously explored for development of targeted therapeutics for malaria and tuberculosis. In this current work, we establish the MEP pathway enzyme DXR as an attractive new therapeutic target for treatment of infections due to zoonotic staphylococci.

The MEP pathway has a number of major advantages as an antimicrobial target for veterinarian applications. Since mammals utilize the mevalonate pathway for isoprenoid biosynthesis, they lack homologs of the MEP pathway enzymes. As a result, MEP pathway inhibition is expected to have a high therapeutic index, and indeed, such inhibitors have been well-tolerated in preclinical and Phase I and II human studies(23– 25,55,56). In addition, use of antibiotics in animal health and agriculture has been implicated as a major driver of antimicrobial resistance in human pathogens(57–60). Of particular relevance to treatment of canine and feline infections, the close physical contact between owners and household pets facilitates not only the cross-colonization of organisms, but also direct transfer of drug-resistance traits(61–63). Because human-associated staphylococci, including *S. aureus, S. warnerii*, and *S. epidermidis*, use the mevalonate pathway for isoprenoid biosynthesis, they are not susceptible to MEP pathway inhibitors (Table 1). Importantly, while Gram-negative organisms such as *E. coli* and *Klebsiella pneumoniae* are modestly susceptible to MEP pathway inhibition, our lipophilic prodrug compound 2 does not inhibit growth of these organisms (Table S5). Our MEPicide compounds therefore have a highly specific and valuable antimicrobial spectrum, which may help break the cycle of resistance transfer from antibiotic-treated animals to the microbiota of humans.

In the current study, we establish the cellular, enzymatic, and structural mechanisms-of-action of FSM against zoonotic staphylococci. We confirm that FSM is a competitive inhibitor of staphylococcal DXR, interrupts cellular isoprenoid biosynthesis, and inhibits growth of zoonotic staphylococci. Of note, the staphylococcal DXR enzyme appears somewhat distinct from previously characterized orthologs, particularly in the α10-α11 loop sequence, which could be explored with additional SAR studies. Together, our work provides insights into differences in staphylococcal DXR that may be key to driving future structure-based inhibitor design efforts.

A well-appreciated liability of antibacterial phosphonates, including fosfomycin and FSM, has been the ready acquisition of resistance through loss of transport (27,64–66). Our work establishes GlpT as the likely phosphonate transporter in zoonotic staphylococci (Fig. 6). Identification of multiple, independent loss-of-function alleles from independent screens in two separate species is compelling evidence for a role of this locus in FSM-resistance in staphylococci. In addition, the homology between staphylococcal GlpT orthologs and Gram-negative phosphonate transporters suggests that the staphylococcal proteins are functionally similar. The finding that lipophilic prodrug MEPicides, which do not require active transport, are still active against the *glpT* mutant strains indicates that the molecular basis of phosphonate resistance is through loss of GlpT-mediated transport (Fig 6). The prodrug MEPicides circumvent GlpT, which our study has shown is easily mutated in staphylococci. Whether staphylococci also readily develop resistance to the prodrug MEPicides is currently unclear, and is an important question for future studies.

It is important to note that while data indicate that the *glpT* mutants are resistant to phosphonate parent compound 3, the magnitude of resistance is substantially less than that of FSM. These data suggest that compound 3 may preferentially use an alternative transporter, thereby bypassing the dependence on GlpT. Surprisingly, staphylococcal *glpT* mutants are hypersensitive to MEPicide prodrugs, suggesting that after penetration and cleavage by cellular esterases, the compounds may accumulate intracellularly in the absence of GlpT (Fig 5). Future studies should examine the cellular transport of the MEPicide compounds, and further, explore whether synergy exists between the parent and prodrug varieties of this class of inhibitors.

The MEPicide prodrugs, including compounds 2 and 4, represent promising leads for ongoing preclinical testing and development of new therapeutics for zoonotic staphylococcal infections. The prodrugs harness the microbial specificity and thus safety of MEP pathway inhibition, while avoiding the dependency on active GlpT-mediated transport. In addition, we find that ester modification has a dramatic effect on anti-staphylococcal potency *in vitro*, suggesting that phosphonate transport limits the anti-bacterial efficacy of FSM and related compounds. Lipophilic ester modifications have previously been employed to improve pharmacokinetic properties and bioavailability of anti-staphylococcal agents (e.g., cefditoren pivoxil)(67). Since MEPicide ester modification at the site of infection is necessary to facilitate bacterial cell entry of inhibitors, future studies will aim to understand what chemical features drive intestinal and serum cleavage of the MEPicide prodrugs.

## Materials and Methods

### DXR Inhibitors

FSM (Millipore Sigma) and FR-900098 (Millipore Sigma) were resuspended in sterile water. Compounds 1-4 were synthesized and resuspended in DMSO as previously described(41,42,53).

### Growth inhibition assays of *Staphylococcus* species

Overnight cultures were diluted 1:200 in LB media and grown at 37°C until the mid-logarithmic phase (OD_600_ = 0.5 – 0.8). Cultures were diluted in a 96-well plate to 1 × 10^5^ in 150 μL LB media and treated with inhibitors at concentrations ranging from 2 nM to 100 μM. Bacteria were grown at 37°C for 20 h with cyclic shaking at 700 rpm in a FLUOstar Omega microplate reader (BMG Labtech). Growth was assessed over 20 h by measuring the OD_600_ at 20 min increments. The half-maximal inhibitory concentration (IC_50_) values were determined during logarithmic growth using GraphPad Prism software. All experiments were performed at least in triplicate and data reported represent the mean ± SEM.

### Minimum bactericidal (MBC) assay

Overnight cultures were diluted 1:200 in LB media and grown at 37°C until reaching mid-logarithmic phase of growth. Compounds were added to cultures at their respective IC_50_ and at 10 × IC_50_, and the bacteria were incubated at 37°C for 24 h while shaking. Cultures were serially diluted in Dulbecco’s Phosphate Buffered Saline (PBS; Gibco) and plated on LB agar. Colonies were enumerated after overnight growth at 37°C. Values reflect the mean and standard deviations of at least three independent experiments.

### Sample preparation for mass spectrometry analysis

Overnight cultures of *Staphylococcus* spp. were diluted 1:200 in LB media and grown at 37°C until reaching mid-logarithmic phase. Cultures were then treated for 2 h with FSM at 10x their IC_50_ while shaking at 37°C. For normalization, the OD_600_ was determined after 2 h of treatment with the DXR inhibitors. Cells were pelleted by centrifugation for 5 min at 3000 × g at 4°C. The supernatants were removed and cells were washed twice with PBS (Gibco). The supernatants were removed and the pellets stored at −80°C until analysis. MEP intermediates were extracted from the samples using glass beads (212-300 u) and 600 μL chilled H_2_O: chloroform: methanol (3:5:12 v/v) spiked with PIPES (piperazine-N,N’-bis(2-ethanesulfonic acid) as internal standard. The cells were disrupted with the TissueLyser II instrument (Qiagen) using a microcentrifuge tube adaptor set pre-chilled for 2 min at 20 Hz. The samples were then centrifuged at 16,000 × g at 4°C for 10 min, the supernatants collected, and pellet extraction repeated once more. The supernatants were pooled and 300 μL chloroform and 450 μL of chilled water were added to the supernatants. The tubes were vortexed and centrifuged. The upper layer was transferred to a 2 mL tube PVDF filter (ThermoFisher, F2520-5) and centrifuged for 5 min at 4,000 × g at 4°C. The samples were transferred to new tubes and dried using a speed-vac. The pellets were re-dissolved in 100 μL of 50% acetonitrile.

### LC-MS/MS analysis

For LC separation, Luna-NH2 column (3 μm, 150 × 2 mm, Phenomenex) was used flowing at 0.4 mL/min. The gradient of the mobile phases A (20 mM ammonium acetate, pH 9.8, 5% acetonitrile) and B (100% acetonitrile) was as follows: 60% B for 1 min, to 6% B in 3 min, hold at 6% B for 5 min, then back to 60% B for 0.5 min. The LC system was interfaced with a Sciex QTRAP 6500^+^ mass spectrometer equipped with a TurboIonSpray (TIS) electrospray ion source. Analyst software (version 1.6.3) was used to control sample acquisition and data analysis. The QTRAP 6500^+^ mass spectrometer was tuned and calibrated according to the manufacturer’s recommendations. The metabolites were detected using MRM transitions that were previously optimized using standards. The instrument was set-up to acquire in negative mode. For quantification, an external standard curve was prepared using a series of standard samples containing different concentrations of metabolites and a fixed concentration of the internal standard. The limit of detection for 1-deoxy-D-xylulose 5-phosphate (DOXP), 4-diphosphocytidyl-2-C-methylerythritol (CDP-ME), and 2-C-methyl-D-erythritol 2,4-cyclopyrophosphate (MEcPP) was 0.0064 μM for a 10 μL injection volume. Data reflect the mean and SD of at least three independent experiments. T-tests were used to test for significance between untreated (UNT) and drug-treated bacteria (Prism).

### Recombinant expression and purification of DXR

Wild-type *dxr* from *S. schleiferi* was amplified from genomic DNA using the forward primer 5’-CTCACCACCACCACCACCAT ATGAAAAATATAGCAATTTTAGGCGC-3’ and the reverse primer 3’-ATCCTATCTTACT CACCTACACCTCATATGATTTTGTTTTATAAT-5’ The PCR product was cloned into vector BG1861 by ligation-independent cloning to introduce a N-terminal 6xHis tag, and transformed into Stellar™ chemically competent cells (Clontech Laboratories)(68). The sequence was confirmed by Sanger sequencing and the plasmid was transformed into *E. coli* BL21(DE3) pLysS (Life Technologies). Gene expression was induced for 2 h with 1 mM isopropyl-β-D-thiogalactoside (IPTG) and cells were harvested by centrifugation at 4274 × g for 10 min at 4°C. The cell pellet was lysed by sonication in lysis buffer containing 25 mM Tris HCl (pH 7.5), 100 mM NaCl, 20 mM imidazole, 10% glycerol, 1 mM MgCl_2_, 1 mM dithiothreitol (DTT), 1 mg/mL lysozyme, 75 U benzonase and 1 Complete Mini EDTA-free protease inhibitor tablet (Roche Applied Science). The hexahistidine-tagged DXR protein was affinity purified from soluble lysate via nickel agarose beads (Gold Biotechnology). Bound protein was eluted in 300 mM imidazole, 25 mM Tris HCl (pH 7.5), 1 mM MgCl_2_ and 100 mM NaCl. Purified protein was dialyzed in buffer containing 10% glycerol without imidazole prior to analysis. The enzyme was frozen in liquid nitrogen and stored permanently at −80°C.

### DXR enzyme activity and inhibitory constant determination

Oxidation of NADPH to NADP^+^ as a result of substrate turnover was monitored at 340 nm in a POLARstar Omega microplate reader (BMG Labtech)(69). The standard reaction had a final concentration of 62.5 nM purified DXR protein, 0.5 mM NADPH, 100 mM NaCl, 25 mM Tris pH 7.5, 10% glycerol, 1 mM MgCl_2_ and 0.09 mg/mL BSA in 50 μL volume per assay. Reactions were initiated by the addition of DOXP after 15 min incubation of the reaction mixture without DOXP at 37°C. Absorption at 340 nm was measured continuously for up to 45 min. For K_m_ [DOXP] determination, DOXP concentrations between 0 and 2 mM were tested at 0.5 mM NADPH. The linear range of enzyme activity was determined by varying the DXR concentration at 1 mM DOXP and 1 mM NADPH. IC_50_ assays were performed using the standard reaction conditions with the respective amount of DXR inhibitor added to obtain the given final concentrations. Data points from at least three independent replicates were analyzed by nonlinear regression using GraphPad Prism software. Slopes of changing absorbance values were converted to (μM DOXP)(mg enzyme)^−1^ s ^−1^ using a NADPH standard curve (data not shown). For the determination of the inhibitory constant Ki [FSM] of DXR, enzyme activity over a range of DOXP substrate concentrations between 0 and 2 mM was measured for FSM between 0 mM to 4 mM. Data points from at least three independent replicates were analyzed as described above.

### Protein crystallography

Crystals of *S. schleiferi* DXR were grown at 4°C using the vapor diffusion method in hanging drops of a 1:1 mixture of protein (10 mg mL^−1^) and crystallization buffer (2 M ammonium sulfate, 100 mM sodium citrate/citric acid, pH 5.5). Crystals of the *S. schleiferi* DXR•FSM complex were obtained in 100 mM HEPES/MOPS (pH 7.5), 20 mM D-glucose, 20 mM D-mannose, 20 mM D-galactose, 20 mM L-fucose, 20 mM D-xylose, 20 mM N-acetyl-D-glucosamine, 20% glycerol, 10% PEG 4000, and 2 mM FSM. Prior to data collection, crystals were stabilized in cryoprotectant (mother liquor supplemented with 30% glycerol) before flash freezing in liquid nitrogen for data collection at 100 K. All diffraction images were collected at beamline 19-ID of the Argonne National Laboratory Advanced Photon Source at Argonne National Laboratory. HKL3000 was used to index, integrate, and scale the data sets(70). For phasing of the apoenzyme structure, molecular replacement was performed in PHASER using the x-ray crystal structure of *E. coli* DXR (PDB: 1T1S) as a search model(31,71). Two monomers were found in the asymmetric unit, with each forming a physiological dimer by crystallographic symmetry. For iterative rounds of model building and refinement, COOTand PHENIX were used, respectively(72,73). The resulting model was used to solve the structure of the FSM complex by molecular replacement with PHASER. Two molecules were found in the asymmetric unit with crystallographic symmetry completing each dimer. Data collection and refinement statistics are summarized in Table S2. Atomic coordinates and structure factors of *S. schleiferi* DXR (PDB:6MH4) and the *S. schleiferi* DXR•FSM complex (PDB:6MH5) were deposited in the RCSB Protein Data Bank.

### Generation of FSM-resistant mutants in *S. schleiferi* and *S. pseudintermedius*

Clinical isolates of *S. schleiferi* (S53022327s) and *S. pseudintermedius* (H20421242p) were cloned and adapted to laboratory media via four rounds of sequential colony isolation and growth on LB agar plates. The isolated FSM-sensitive parental clones were incubated overnight on LB agar containing FSM (32 μM). Surviving single colonies were re-struck onto LB agar for clonal isolation. FSM resistance of isolated clones was confirmed by overnight growth on LB agar containing FSM (32 μM). The FSM-sensitive parental clones were used as a control to confirm growth and antibiotic-resistance.

### Quantification of MEPicide resistance

Minimum Inhibitory Concentration (MIC) assays were conducted by microtiter broth dilution in clear 96-well plates(74). MEPicides were serially diluted in duplicate at concentrations ranging from 1.5 mM – 19.5 nM in 75 μL of LB broth. Bacteria cultured without drug were used as a positive control for growth. The plates were inoculated with 75 μL bacteria diluted to 1 × 10^5^ CFU/mL in LB. Plates were incubated for 18-20 h while shaking at 200 RPM at 37°C. The plates were then visually inspected, and the MIC value was defined as the lowest concentration of MEPicide that prevented visual growth.

### Whole genome sequencing and variant discovery

Genomic DNA was isolated from overnight cultures of *S. pseudintermedius* and *S. schleiferi* using a standard phenol-chloroform extraction and ethanol precipitation protocol. Sequencing libraries were prepared and sequenced by the Washington University Genome Technology Access Center (GTAC). 1 μg of DNA was sonicated to an average size of 175 bp. Fragments were blunt ended and had an A base added to the 3’ end. Sequence adapters were ligated to the ends and the sequence tags were added via amplification. Resulting libraries were sequenced on an Illumina HiSeq 2500 to generate 101 bp paired end reads. DNA quantity and quality were assessed by GTAC using Agilent Tapestation.

For WGS, sequences from GenBank were retrieved from the following organisms: *S. pseudintermedius* ED99 (accession number CP002478) and *S. schleiferi* 1360-13 (CP009740) assemblies were downloaded from NCBI (ftp://ftp.ncbi.nlm.nih.gov). Paired-end reads were aligned to each of the available genomes using Novoalign v3.03. (Novocraft Technologies) and deposited in NCBI (accession number PRJNA488092). Duplicates were removed and variants were called using SAMtools(75). SNPs were filtered against parent variants and by mean depth value and quality score (minDP =5, minQ = 37)(76). Genetic variants were annotated using SnpEff v4.3 (Table S4)(77). For all samples, at least 90% of the genome was sequenced at 20x coverage. All whole genome sequencing data is available in the NCBI BioProject database and Sequence Read Archive. Point mutations found in the GlpT domain were mapped onto the predicted transmembrane topology of GlpT using Protter(78).

### Sanger Sequencing of *S. schleiferi* and *S. pseudintermedius glpT*

Reference sequences for *glpT* in *S. schleiferi* (<u>WP_016426432.1</u>) and *S. pseudintermedius* (<u>WP_014613322.1</u>) were found with the Basic Local Alignment Search Tool (BLAST, v. 2/2/22). The regions of interest were amplified from *S. pseudintermedius* and *S. schleiferi* using gene-specific primers (Table S1). Amplicons were sequenced by the Washington University Protein and Nucleic Acid Laboratory using BigDye Terminator v3.1 Cycle Sequencing reagents (Life Technologies). Representative traces for all strains are available through the NCBI Trace Archive.

## Acknowledgements

The authors are grateful to David Hunstad, Timothy Hagen, and Phillip Tarr for providing bacterial strains used in these studies. We would also like to thank Meghan Wallace for assistance with MALDI-TOF mass spectrometry identification of the Staphylococcal isolates.

## Supporting Information

**S1 Fig. DXR inhibitors are bacteriostatic.** Growth in CFU/mL of *S. schleiferi* and *S. pseudintermedius* after 24 h treatment is plotted against the respective treatment. Cultures were treated at 1 × IC_50_ concentration and/or 10x IC_50_ concentration of the inhibitors. Shown are the mean values + SD from at least three independent experiments.

**S2 Fig. SDS-PAGE of purified *S. schleiferi* DXR**. Molecular mass standard (M) and approximately 1 μg of purified recombinant *S. schleiferi* DXR.

**S3 Fig. Membrane topology of GlpT.** (A) Wild-type amino acid sequences and predicted transmembrane topology of *S. schleiferi* GlpT. Residues Gly-99, Trp-148, Trp-161, Ala-267, Gly-298, Ala-309, and Gln-379 are indicated in the sequence. Red indicates a stop mutation at the site, while blue indicates a missense mutation. (B) Wild-type amino acid sequences and predicted transmembrane topology of *S. pseudintermedius* GlpT. Residues Asp-88, Gly-99, Gly-135, Trp-301, Gly-400, and Gly-404 are indicated in the sequence. Red indicates a stop mutation at the site, while blue indicates a missense mutation. Schematic diagrams were prepared with the program Protter(5).

**S1 Table. Primers.**

**S2 Table. Summary of crystallographic data collection and refinement statistics.**

**S3. Table. FSM MICs, *glpT* alleles, GlpT protein changes, and Polyphen-2 scores for FSM**^**R**^ **strains.**

**S4. Table. SNP calls from FSM**^**R**^ ***S. schleiferi* and *S. pseudintermedius* strains.** Genomes were aligned to reference genomes *S. schleiferi* 1360-13 and *S. pseudintermedius* ED99, respectively. Each line represents a SNP call. Changes shown are those not present in the parental strain. Changes determined to be false by Sanger sequencing have been removed. GlpT is highlighted in green. ^*^Location of the change inside the gene, ^†^the base at that location, ^‡^the new base present at that location, ^§^the corresponding protein change associated with the new base, ^¶^the gene name according to the previous annotation, ^#^the predicted function.

**S5 Table. Inhibitory effect of MEPicides against a panel of Gram-negative bacteria.** IC_50_ values are reported in μM. Data represent the mean ± SD from at least three independent experiments.

**S1. File. Supplemental Methods.**

## References

1. Jindal A, Shivpuri D, Sood S. Staphylococcus schleiferi meningitis in a child. Pediatr Infect J. 2015;34(3):329.

2. Somayaji R, Rubin JE, Priyantha MA, Church D. Exploring *Staphylococcus pseudintermedius:* an emerging zoonotic pathogen? Future Microbiol. 2016;11(11):1371–4.

3. Börjesson S, Gómez-Sanz E, Ekström K, Torres C, Grönlund U. *Staphylococcus pseudintermedius* can be misdiagnosed as *Staphylococcus aureus* in humans with dog bite wounds. Eur J Clin Microbiol Infect Dis. 2015;34(4):839–44.

4. Lainhart W, Yarbrough ML, Burnham CA. The brief case: *Staphylococcus intermedius* group-look what the dog dragged in. J Clin Microbiol. 2018;56(2).

5. Rojas-Marte G, Victor J, Shenoy A, Yakubov S, Chapnick E, Lin YS. Pacemaker-associated infective endocarditis caused by *Staphylococcus schleiferi*. Infect Dis Clin Pract. 2014;22(5):302–4.

6. Pottumarthy S, Schapiro JM, Prentice JL, Houze YB, Swanzy SR, Fang FC, et al. Clinical isolates of *Staphylococcus intermedius* masquerading as methicillinresistant *Staphylococcus aureus*. J Clin Microbiol. 2004;42(12):5881–4.

7. Yarbrough ML, Lainhart W, Burnham CA. Epidemiology, clinical characteristics, and antimicrobial susceptibility profiles of human clinical isolates of *Staphylococcus intermedius* group. J Clin Microbiol. 2018;56(3):e01788–17.

8. Ross Fitzgerald J. The *Staphylococcus intermedius* group of bacterial pathogens: species re-classification, pathogenesis and the emergence of meticillin resistance. Vet Dermatol. 2009;20(5–6):490–5.

9. Humphries RM, Wu MT, Westblade LF, Robertson AE, Burnham C-AD, Wallace MA, et al. In vitro antimicrobial susceptibility of Staphylococcus pseudintermedius isolates of human and animal origin. J Clin Microbiol. 2016;54(5):1391–4.

10. Beever L, Bond R, Graham PA, Jackson B, Lloyd DH, Loeffler A. Increasing antimicrobial resistance in clinical isolates of *Staphylococcus intermedius* group bacteria and emergence of MRSP in the UK. Vet Rec. 2015;176(7):172.

11. Lange BM, Rujan T, Martin W, Croteau R. Isoprenoid biosynthesis: The evolution of two ancient and distinct pathways across genomes. Proc Natl Acad Sci. 2000;97(24):13172–7.

12. Wilding EI, Kim DY, Bryant AP, Gwynn MN, Lunsford RD, McDevitt D, et al. Essentiality, expression, and characterization of the class II 3-hydroxy-3-methylglutaryl coenzyme A reductase of *Staphylococcus aureus*. J Bacteriol. 2000;182(18):5147–52.

13. Matsumoto Y, Yasukawa J, Ishii M, Hayashi Y, Miyazaki S, Sekimizu K. A critical role of mevalonate for peptidoglycan synthesis in *Staphylococcus aureus*. Sci Rep. 2016;6:22894.

14. Liu C-I, Liu GY, Song Y, Yin F, Hensler ME, Jeng W-Y, et al. A cholesterol biosynthesis inhibitor blocks *Staphylococcus aureus* virulence. Science. 2008;319(5868):1391–4.

15. Misic AM, Cain CL, Morris DO, Rankin SC, Beiting DP. Divergent isoprenoid biosynthesis pathways in *Staphylococcus* species constitute a drug target for treating infections in companion animals. mSphere. 2014;1(5):1–11.

16. Nair SC, Brooks CF, Goodman CD, Sturm A, Strurm A, McFadden GI, et al. Apicoplast isoprenoid precursor synthesis and the molecular basis of fosmidomycin resistance in *Toxoplasma gondii*. J Exp Med. 2011;208(7):1547– 59.

17. Odom AR, Van Voorhis WC. Functional genetic analysis of the *Plasmodium falciparum* deoxyxylulose 5-phosphate reductoisomerase gene. Mol Biochem Parasitol. 2010;170(2):108–11.

18. Brown AC, Parish T. Dxr is essential in *Mycobacterium tuberculosis* and fosmidomycin resistance is due to a lack of uptake. BMC Microbiol. 2008;8:78.

19. McAteer S, Coulson A, McLennan N, Masters M. The *lytB* gene of *Escherichia coli* is essential and specifies a product needed for isoprenoid biosynthesis. J Bacteriol. 2001;183(24):7403–7.

20. Wagner WP, Helmig D, Fall R. Isoprene biosynthesis in Bacillus subtilis via the methylerythritol phosphate pathway. 1999; doi:10.1021/NP990286P.

21. McKenney ES, Sargent M, Khan H, Uh E, Jackson ER, San Jose G, et al. Lipophilic prodrugs of FR900098 are antimicrobial against *Francisella novicida in vivo* and *in vitro* and show GlpT independent efficacy. PLoS One. 2012;7(10):e38167.

22. Koppisch AT, Fox DT, Blagg BSJ, Poulter CD. E. coli MEP synthase: Steadystate kinetic analysis and substrate binding. Biochemistry. 2002;41(1):236–43.

23. Kuemmerle HP, Murakawa T, Sakamoto H, Sato N, Konishi T, De Santis F. Fosmidomycin, a new phosphonic acid antibiotic. Part II: 1. Human pharmacokinetics. 2. Preliminary early phase IIa clinical studies. Int J Clin Pharmacol Ther Toxicol. 1985;23(10):521–8.

24. Borrmann S, Lundgren I, Oyakhirome S, Impouma B, Matsiegui P-B, Adegnika AA, et al. Fosmidomycin plus clindamycin for treatment of pediatric patients aged 1 to 14 years with *Plasmodium falciparum* malaria. Antimicrob Agents Chemother. 2006;50(8):2713–8.

25. Tsuchiya T, Ishibashi K, Terakawa M, Nishiyama M, Itoh N, Noguchi H. Pharmacokinetics and metabolism of fosmidomycin, a new phosphonic acid, in rats and dogs. Eur J Drug Metab Pharmacokinet. 7(1):59–64.

26. Dhiman RK, Schaeffer ML, Bailey AM, Testa CA, Scherman H, Crick DC. 1-deoxy-D-xylulose 5-phosphate reductoisomerase (IspC) from *Mycobacterium tuberculosis*: towards understanding mycobacterial resistance to fosmidomycin. J Bacteriol. 2005;187(24):8395–402.

27. Mackie RS, McKenney ES, van Hoek ML. Resistance of *Francisella novicida* to fosmidomycin associated with mutations in the glycerol-3-phosphate transporter. Front Microbiol. 2012;3:226.

28. Sakamoto Y, Furukawa S, Ogihara H, Yamasaki M. Fosmidomycin resistance in adenylate cyclase deficient (*cya*) mutants of *Escherichia coli*. Biosci Biotechnol Biochem. 2003;67(9):2030–3.

29. Argyrou A, Blanchard JS. Kinetic and chemical mechanism of Mycobacterium tuberculosis 1-deoxy-D-xylulose-5-phosphate isomeroreductase. Biochemistry. 2004;43(14):4375–84.

30. Kuzuyama T, Takahashi S, Takagi M, Seto H. Characterization of 1-deoxy-D-xylulose 5-phosphate reductoisomerase, an enzyme involved in isopentenyl diphosphate biosynthesis, and identification of its catalytic amino acid residues. J Biol Chem. 2000;275(26):19928–32.

31. Shunsuke Yajima, Kodai Hara, John M. Sanders, Fenglin Yin, Kanju Ohsawa, Jochen Wiesner, et al. Crystallographic structures of two bisphosphonate:1-deoxyxylulose-5-phosphate reductoisomerase complexes. 2004; doi:10.1021/JA040126M.

32. Yajima S, Nonaka T, Kuzuyama T, Seto H, Ohsawa K. Crystal structure of 1-deoxy-D-xylulose 5-phosphate reductoisomerase complexed with cofactors: implications of a flexible loop movement upon substrate binding. J Biochem. 2002;131(3):313–7.

33. Yajima S, Hara K, Iino D, Sasaki Y, Kuzuyama T, Ohsawa K, et al. Structure of 1-deoxy-D-xylulose 5-phosphate reductoisomerase in a quaternary complex with a magnesium ion, NADPH and the antimalarial drug fosmidomycin. Acta Crystallogr Sect F Struct Biol Cryst Commun. 2007;63(Pt 6):466–70.

34. Behrendt CT, Kunfermann A, Illarionova V, Matheeussen A, Pein MK, Gräwert T, et al. Reverse fosmidomycin derivatives against the antimalarial drug target IspC (Dxr). J Med Chem. 2011;54(19):6796–802.

35. Deng L, Endo K, Kato M, Cheng G, Yajima S, Song Y. Structures of 1-deoxy-D-xylulose-5-phosphate reductoisomerase/lipophilic phosphonate complexes. ACS Med Chem Lett. 2011;2(2):165–70.

36. Sooriyaarachchi S, Chofor R, Risseeuw MDP, Bergfors T, Pouyez J, Dowd CS, et al. Targeting an aromatic hotspot in *Plasmodium falciparum* 1-deoxy-D-xylulose-5-phosphate reductoisomerase with β-arylpropyl analogues of fosmidomycin. ChemMedChem. 2016;11(18):2024–36.

37. Mac Sweeney A, Lange R, Fernandes RPM, Schulz H, Dale GE, Douangamath A, et al. The crystal structure of *E. coli* 1-deoxy-D-xylulose-5-phosphate reductoisomerase in a ternary complex with the antimalarial compound fosmidomycin and NADPH reveals a tight-binding closed enzyme conformation. J Mol Biol. 2005;345(1):115–27.

38. Kholodar SA, Tombline G, Liu J, Tan Z, Allen CL, Gulick AM, et al. Alteration of the flexible loop in 1-deoxy-D-xylulose-5-phosphate reductoisomerase boosts enthalpy-driven inhibition by fosmidomycin. Biochemistry. 2014;53(21):3423–31.

39. Adzhubei IA, Schmidt S, Peshkin L, Ramensky VE, Gerasimova A, Bork P, et al. A method and server for predicting damaging missense mutations. Nat Methods. 2010;7(4):248–9.

40. Murakawa T, Sakamoto H, Fukada S, Konishi T, Nishida M. Pharmacokinetics of fosmidomycin, a new phosphonic acid antibiotic. Antimicrob Agents Chemother. 1982;21(2):224–30.

41. Jackson ER, San Jose G, Brothers RC, Edelstein EK, Sheldon Z, Haymond A, et al. The effect of chain length and unsaturation on Mtb Dxr inhibition and antitubercular killing activity of FR900098 analogs. Bioorg Med Chem Lett. 2014;24(2):649–53.

42. Edwards RL, Brothers RC, Wang X, Maron MI, Ziniel PD, Tsang PS, et al. MEPicides: potent antimalarial prodrugs targeting isoprenoid biosynthesis. Sci Rep. 2017;7(1):8400.

43. Uh E, Jackson ER, San Jose G, Maddox M, Lee RE, Lee RE, et al. Antibacterial and antitubercular activity of fosmidomycin, FR900098, and their lipophilic analogs. Bioorg Med Chem Lett. 2011;21(23):6973–6.

44. San Jose G, Jackson ER, Haymond A, Johny C, Edwards RL, Wang X, et al. Structure–Activity Relationships of the MEPicides: *N* -Acyl and *O* -Linked Analogs of FR900098 as Inhibitors of Dxr from *Mycobacterium tuberculosis* and *Yersinia pestis*. ACS Infect Dis. 2016;2(12):923–35.

45. Brücher K, Gräwert T, Konzuch S, Held J, Lienau C, Behrendt C, et al. Prodrugs of reverse fosmidomycin analogues. J Med Chem. 2015;58(4):2025–35.

46. Faísca Phillips AM, Nogueira F, Murtinheira F, Barros MT. Synthesis and antimalarial evaluation of prodrugs of novel fosmidomycin analogues. Bioorg Med Chem Lett. 2015;25(10):2112–6.

47. Brücher K, Illarionov B, Held J, Tschan S, Kunfermann A, Pein MK, et al. α-Substituted β-oxa isosteres of fosmidomycin: synthesis and biological evaluation. J Med Chem. 2012;55(14):6566–75.

48. Haemers T, Wiesner J, Giessmann D, Verbrugghen T, Hillaert U, Ortmann R, et al. Synthesis of beta-and gamma-oxa isosteres of fosmidomycin and FR900098 as antimalarial candidates. Bioorg Med Chem. 2008;16(6):3361–71.

49. Wiesner J, Ortmann R, Jomaa H, Schlitzer M. Double ester prodrugs of FR900098 display enhanced *in-vitro* antimalarial activity. Arch Pharm (Weinheim). 2007;340(12):667–9.

50. Kurz T, Schlüter K, Pein M, Behrendt C, Bergmann B, Walter RD. Conformationally restrained aromatic analogues of fosmidomycin and FR900098. Arch Pharm (Weinheim). 2007;340(7):339–44.

51. Kurz T, Schlüter K, Kaula U, Bergmann B, Walter RD, Geffken D. Synthesis and antimalarial activity of chain substituted pivaloyloxymethyl ester analogues of fosmidomycin and FR900098. Bioorg Med Chem. 2006;14(15):5121–35.

52. Schlüter K, Walter RD, Bergmann B, Kurz T. Arylmethyl substituted derivatives of fosmidomycin: synthesis and antimalarial activity. Eur J Med Chem. 2006;41(12):1385–97.

53. Wang X, Edwards R, Ball H, Johnson C, Haymond A, Girma M, et al. MEPicides: α,β-unsaturated fosmidomycin analogs as DXR inhibitors against malaria. J Med Chem. 2018;61(19):8847–58.

54. Weese JS, van Duijkeren E. Methicillin-resistant *Staphylococcus aureus* and *Staphylococcus pseudintermedius* in veterinary medicine. Vet Microbiol. 2010;140(3–4):418–29.

55. Kuemmerle HP, Murakawa T, Soneoka K, Konishi T. Fosmidomycin: a new phosphonic acid antibiotic. Part I: Phase I tolerance studies. Int J Clin Pharmacol Ther Toxicol. 1985;23(10):5151–20.

56. Borrmann S, Adegnika AA, Moussavou F, Oyakhirome S, Esser G, Matsiegui P-B, et al. Short-course regimens of artesunate-fosmidomycin in treatment of uncomplicated *Plasmodium falciparum* malaria. Antimicrob Agents Chemother. 2005;49(9):3749–54.

57. Silbergeld EK, Graham J, Price LB. Industrial food animal production, antimicrobial resistance, and human health. Annu Rev Public Health. 2008;29(1):151–69.

58. Holmes A, Moore L, Sundsfjord A, Steinbakk M, Regmi S, Karkey A, et al. Understanding the mechanisms and drivers of antimicrobial resistance. Lancet. 2016;387(10014):176–87.

59. Robinson TP, Bu DP, Carrique-Mas J, Fèvre EM, Gilbert M, Grace D, et al. Antibiotic resistance is the quintessential One Health issue. Trans R Soc Trop Med Hyg. 2016;110(7):377–80.

60. Robinson TP, Wertheim HFL, Kakkar M, Kariuki S, Bu D, Price LB. Animal production and antimicrobial resistance in the clinic. Lancet. 2016;387(10014):e1–3.

61. Guardabassi L, Schwarz S, Lloyd DH. Pet animals as reservoirs of antimicrobial-resistant bacteria. Vol. 54, Journal of Antimicrobial Chemotherapy. 2004. p. 321– 32.

62. Martins LRL, Pina SMR, Simões RLR, de Matos AJF, Rodrigues P, da Costa PMR. Common phenotypic and genotypic antimicrobial resistance patterns found in a case study of multiresistant E. coli from cohabitant pets, humans, and household surfaces. J Environ Health. 75(6):74–81.

63. Lloyd DH. Reservoirs of antimicrobial resistance in pet animals. Clin Infect Dis. 2007 Sep 1;45(Supplement_2):S148–52.

64. Takahata S, Ida T, Hiraishi T, Sakakibara S, Maebashi K, Terada S, et al. Molecular mechanisms of fosfomycin resistance in clinical isolates of *Escherichia coli*. Int J Antimicrob Agents. 2010;35(4):333–7.

65. Chekan JR, Cogan DP, Nair SK. Molecular basis for resistance against phosphonate antibiotics and herbicides. Medchemcomm. 2016;7(1):28–36.

66. Castañeda-García A, Blázquez J, Rodríguez-Rojas A. Molecular mechanisms and clinical impact of acquired and intrinsic fosfomycin resistance. Antibiot (Basel, Switzerland). 2013;2(2):217–36.

67. Guay DRP. Review of cefditoren, an advanced-generation, broad-spectrum oral cephalosporin. Clin Ther. 2001;23(12):1924–37.

68. Alexandrov A, Vignali M, LaCount DJ, Quartley E, de Vries C, De Rosa D, et al. A facile method for high-throughput co-expression of protein pairs. Mol Cell Proteomics. 2004;3(9):934–8.

69. Armstrong CM, Meyers DJ, Imlay LS, Freel Meyers C, Odom AR. Resistance to the antimicrobial agent fosmidomycin and an FR900098 prodrug through mutations in the deoxyxylulose phosphate reductoisomerase gene (dxr). Antimicrob Agents Chemother. 2015;59(9):5511–9.

70. Minor W, Cymborowski M, Otwinowski Z, Chruszcz M, IUCr. *HKL* -3000: the integration of data reduction and structure solution – from diffraction images to an initial model in minutes. Acta Crystallogr Sect D Biol Crystallogr. 2006;62(8):859–66.

71. McCoy AJ, Grosse-Kunstleve RW, Adams PD, Winn MD, Storoni LC, Read RJ. Phaser crystallographic software. J Appl Crystallogr. 2007;40(Pt 4):658–74.

72. Emsley P, Cowtan K, IUCr. *Coot*: model-building tools for molecular graphics. Acta Crystallogr Sect D Biol Crystallogr. 2004;60(12):2126–32.

73. Adams PD, Afonine P V., Bunkóczi G, Chen VB, Davis IW, Echols N, et al. *PHENIX*?: a comprehensive Python-based system for macromolecular structure solution. Acta Crystallogr Sect D Biol Crystallogr. 2010;66(2):213–21.

74. Determination of minimum inhibitory concentrations (MICs) of antibacterial agents by broth dilution. Clin Microbiol Infect. 2003;9(8):ix–xv.

75. Li H, Handsaker B, Wysoker A, Fennell T, Ruan J, Homer N, et al. The sequence alignment/map format and SAMtools. Bioinformatics. 2009;25(16):2078–9.

76. Danecek P, Auton A, Abecasis G, Albers CA, Banks E, DePristo MA, et al. The variant call format and VCFtools. Bioinformatics. 2011;27(15):2156–8.

77. Cingolani P, Platts A, Wang LL, Coon M, Nguyen T, Wang L, et al. A program for annotating and predicting the effects of single nucleotide polymorphisms, SnpEff: SNPs in the genome of *Drosophila melanogaster* strain *w1118; iso-2; iso-3*. Fly (Austin). 2012;6(2):80–92.

78. Omasits U, Ahrens CH, Müller S, Wollscheid B. Protter: interactive protein feature visualization and integration with experimental proteomic data. Bioinformatics. 2014;30(6):884–6.

